# Excitatory effect of biphasic kHz field stimulation on CA1 pyramidal neurons in slices

**DOI:** 10.1101/2021.12.26.474217

**Authors:** Sergei Karnup, William De Groat, Jonathan Beckel, Changfeng Tai

## Abstract

**Background:** Electrical stimulation in the kilohertz-frequency range has been successfully used for treatment of various neurological disorders. Nevertheless, the mechanisms underlying this stimulation are poorly understood.

**Objective:** To study the effect of kilohertz-frequency electric fields on neuronal membrane biophysics we developed a reliable experimental method to measure responses of single neurons to kilohertz field stimulation in brain slice preparations.

**Methods:** In the submerged brain slice pyramidal neurons of the CA1 subfield were recorded in the whole-cell configuration before, during and after stimulation with an external electric field at 2kHz, 5kHz or 10 kHz.

**Results:** Reproducible excitatory changes in rheobase and spontaneous firing were elicited during kHz-field application at all stimulating frequencies. The rheobase only decreased and spontaneous firing either was initiated in silent neurons or became more intense in previously spontaneously active neurons. Response thresholds were higher at higher frequencies. Blockade of glutamatergic synaptic transmission did not alter the magnitude of responses. Inhibitory synaptic input was not changed by kilohertz field stimulation.

**Conclusion:** kHz-frequency current applied in brain tissue has an excitatory effect on pyramidal neurons during stimulation. This effect is more prominent and occurs at a lower stimulus intensity at a frequency of 2kHz as compared to 5kHz and 10kHz.

## Introduction

Electrical stimulation of the nervous system is an established treatment for a wide range of neurological disorders. Low frequency electrical fields within the physiological range, i.e. <200 Hz are used clinically for targeted transcranial stimulation of deep brain structures [1] to elicit paresthesia-free pain relief [2, 3], improved control of motor responses, to restore tactile perception [4],and to induce pseudo-spontaneous activity mimicking physiological conditions [5]. The proposed mechanisms for conventional electrical stimulation of < 500 Hz include one-to-one entrainment [6], direct presynaptic depression [7], direct activation or inhibition of neurons [7], and antidromic or orthodromic neuronal excitation [8]. Despite the clinical efficacy of deep brain stimulation (DBS, frequencies <200 Hz) for treatment of Parkinson’s disease, dystonia and tremor, the mechanisms underlying its beneficial effects are poorly understood and are still under debate. Currently there are at least three main hypotheses claiming to explain DBS action on the brain: (1) inhibition hypothesis suggesting that DBS inhibits local neuronal elements, (2) excitation hypothesis suggesting that DBS excites local neurons, and (3) disruption hypothesis suggesting that DBS disrupts abnormal information flow [9]. Such controversy is probably due to consideration of results obtained with different methods in different brain regions with different network architecture and different neuronal properties.

Recently neuromodulation techniques employing kilohertz frequency field stimulation (kHz-FS) have also been used for transcranial brain stimulation [10, 11], spinal cord stimulation [12, 13] and deep brain stimulation [14, 15]. Physiological responses to application of kHz frequencies can occur in a form of facilitation [16], desynchronization [5], axonal conduction block [17, 18] or synaptic fatigue [19]. However, the mechanisms underlying the action kHz-FS at the level of cellular membrane properties and impact on membrane ion currents remain unclear. Thus, to study the interaction between kHz-FS with the neuronal membrane one needs to monitor a single cell isolated (pharmacologically or mechanistically) from its network before, during and after stimulation. However, reproducible reactions of individual neurons to kHz-FS are not available in the literature.

There are several reasons obstructing and discouraging experimental studies of kHz-FS effects. First, low-pass characteristics of neuronal membranes do not allow for direct registration of depolarizing-hyperpolarizing membrane currents during application of kilohertz frequencies [20-22]. Second, in the reported current-clamp whole-cell recordings during kHz-field application in a slice the intracellular voltage was masked by a strong high-frequency noise and an artifactual hyperpolarizing shift was observed due to the absence of the voltage follower in the headstage [23]. Third, measurements of population postsynaptic activity in the hippocampal slice before and after short (1s) or long (up to 30 min) kHz-frequency electric field application did not reveal significant effects either immediately or 30 and 60 min after termination of stimulation [24]. This is in contrast with the long lasting effects of kHz stimulation to produce peripheral axonal conduction block [25-27], or to produce central nervous system changes after spinal cord stimulation [28] and deep brain stimulation in *in vivo* experiments [14]. This inconsistency might be explained either by different mechanisms of action *in vivo* and *in vitro*, or by differences in methodological approach.

## Methods

In this study C57BL/6J mice P60-P120 of either sex were used. Whole cell recordings of hippocampal CA1 pyramidal neurons were made in brain slices submerged in the artificial cerebrospinal fluid (ACSF). To obtain brain slices a mouse was anesthetized with isoflurane and decapitated. After removing the upper part of the cranium, the brain was placed for 5 min in ice-cold sucrose-based saline equilibrated with carbogen (95% O2+5% CO2) and containing: (in mM) 26 NaHCO3, 1 NaH2PO4, 3 KCl, 11 glucose, 234 sucrose, 10 MgSO4, 0.5 CaCl2. Then the whole brain was glued to the vibratome stage with an agar supporting block and cut in to 350 µm thick transverse slices. After cutting, slices were warmed at 32°-33°C for 0.5 hour and then incubated during the experiment at room temperature (23°-24°C) in ACSF saturated with carbogen: (in mM) 117 NaCl, 26 NaHCO3, 3.6 KCl, 1.25 NaH2PO4, 2 MgSO4, 2 CaCl2, and 11 glucose; 285–290 mOsm, pH 7.4.

For a whole cell recording, a single slice was placed in the chamber (∼1 ml volume) and continuously perfused with ACSF at the rate 3–5 ml/min. All recordings were performed at room temperature. The recording chamber was installed on an upright microscope (Olympus BX51W1, Life Science Solutions, Global) equipped for epifluorescence and near-infrared differential interference contrast (DIC) optics. A CCD Hamamatsu C10600 ORCA-R2 camera (Hamamatsu Photonics K.K., Japan) and MetaMorph software package (Molecular Devices, Sunnyvale, CA) were used for visualization of the CA1 pyramidal cell layer of the hippocampus and placement of stimulating electrodes. Patch pipettes were pulled from borosilicate glass capillaries with an inner filament (1.5 mm outer diameter; World Precision Instruments Inc., Sarasota, FL) on a pipette puller (P-97; Sutter Instruments, Novato, CA) and were filled with a solution of the following composition (in mM): 114 K-gluconate, 5 KCl, 0.5 CaCl2, 2 MgCl2, 5 HEPES, 5 EGTA. Osmolarity was adjusted to 270–275 mOsm, pH 7.3. The pipette resistance was 10–14 MΩ. Pyramidal neurons of the CA1 subfield were found blindly in voltage-clamp mode at a depth 70-100 µm under the upper surface of a slice. Recordings were made in the current-clamp mode with the MultiClamp 700B amplifier (Molecular Devices, Sunnyvale, CA) and data acquisition software package Signal5 (Cambridge Electronic Design Limited, Cambridge, UK). There was no correction for the liquid junction potential (∼10 mV).

Bipolar stimulating electrodes were made from 100 µm thick platinum-iridium wire insulated with lacquer. Two wires were inserted into a glass pipette and fixed with CrazyGlue leaving ∼10 mm of their distal ends free. These ends were split to be 1 mm apart, and slightly bent at an angle to make the last 2 mm parallel to the slice (Fig.1). Insulation was removed only from the lower surface (between 5 and 7 o’clock in the cross-section) of each 2 mm long wire ends. Before beginning an experiment these bare surfaces were placed along the CA1 subfield, so that the pyramidal cell layer, *stratum lacunosum* and *stratum oriens* were between the wires, but the CA3 subfield was not covered by an imaginary rectangle of the isopotential field (Fig.1 upper panel). The electrodes were pressed 20-30 µm deep into the slice (Fig.1, lower panel). We recorded neurons located near the midline between these 2 mm long wires. Since the portion of the slice between parallel wires was exposed to an isopotential electric field during electrical stimulation, a precise location of a recorded neuron was not important provided it was close to the middle of the wires’ length and the middle of the distance between the wires. To test the effect of partial insulation we tried to apply kHz electric field through bare wires of a slightly larger length (5 mm). When electrodes were not insulated and were only touching the slice, the effect of the field was substantially weaker. As shown in Figure 2, 3 mA current at 3 different frequencies using uninsulated electrodes did not elicit detectable changes in the electrical properties of the neurons, although with the partially insulated electrodes used in this study 3 mA current elicited changes at all frequencies. Biphasic electric current trains of 2kHz, 5kHz and 10 kHz were generated by custom made script and applied to the slice through a stimulus isolator A395 (WPI, Lawrence, PA). Occasional DC offsets were monitored and neutralized when they occurred. Positive and negative phases of the square waves had equal amplitudes without any gap between phases. Amplitude of biphasic stimulating currents was measured from zero to negative or positive maximum amplitude. All recordings consisted of 10 min long sweeps sequentially stitched during off-line analysis. Ten-to-twenty minutes after membrane rupture and relative stabilization of Vm the baseline activity and membrane properties were measured for 5-10 min; recordings began after this initial stabilization. Trains of extracellular kHz-field stimulation (kHz-FS) lasted for 300 or 600 sec and were intermingled with quiescent periods (200-600 sec) without extracellular field stimulation. We could observe meaningful patterns of neuronal activity during kHz-FS only if the sampling frequency was equal to the stimulation frequency (Fig.2A), whereas if the two frequencies differed, the recording was masked by a high-frequency noise (Fig.2 B, C). Even a slight mismatch between stimulating and sampling frequencies resulted in a high frequency noise. We assumed that high frequency noise recorded during stimulation may result from inequality of sampling frequency and stimulating frequency. Indeed, equalizing the two frequencies resulted in a clearly recorded intracellular signal superimposed on a slow wave; which could be subtracted later during analysis. Fig.3 schematically describes the occurrence of a slow wave when the sampling and stimulation frequencies are equal but phase-shifted (Fig.3A) or have a subtle discrepancy in frequency (Fig.3B), or are substantially different (Fig.3C). When they were equal and the high-frequency noise was absent during kHz stimulation, we still could not measure Vm values due to strong distortion of the recording by an overlapping slow wave (Fig.2 A, 3B). The moments of turning kHz-FS on and off were distinctly separated from periods without stimulation by occurrence and disappearance of this slow wave overlapped on the intracellular signal (Fig.4). A period of this slow oscillation negatively correlated with stimulation frequency: 239 sec for 2 kHz (Fig.5A), 86 sec for 5 kHz (Fig. 4) and 46 sec for 10 kHz (Fig.5B). The amplitude of a slow wave positively correlated with the amplitude of the stimulating current and reversely correlated with input resistance of a cell. The slow wave was seemingly due to a tiny difference between the two nominally equal frequencies because the stimulating train was generated by one computer, and sampling frequency was defined by the other computer. Therefore, each next data point was collected with a tiny phase shift on the stimulating curve. Subtraction of this slow wave from the original record using custom made script allowed measurements of Rin, Rh and, if present, spontaneous firing rate throughout the experiment (Fig.5C). However, direct measurement of Vm during kHz-FS was not possible.

**Fig. 1.**
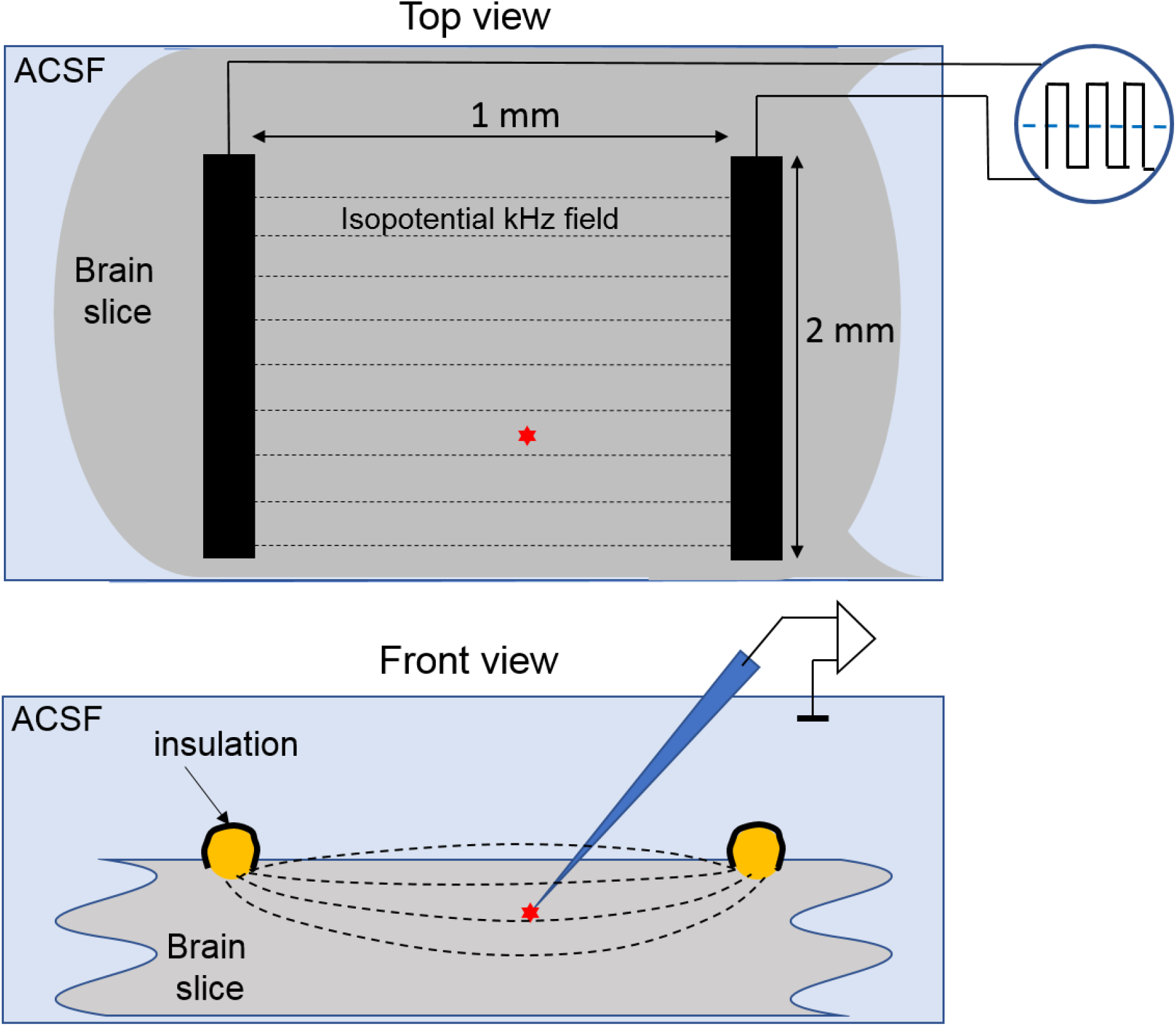
Positioning of electrodes for extracellular field stimulation in the submerged slice. On the upper panel (top view) two black rectangles represent platinum-iridium wires parallel to each other and to the slice surface in the horizontal plane. They were insulated with the lacquer (black cover on the side and top surfaces as seen on cross-sections on the lower panel) except for the bottom surface to minimize shunting of applied currents by surrounding ACSF and to maximize current through the tissue. The recorded neuron (red asterisk) is located between the bars close to the middle of their length. Before the experiment the wire bars were pressed into the slice to a depth of ∼30-50 µm as shown in the lower panel.

**Fig. 2.**
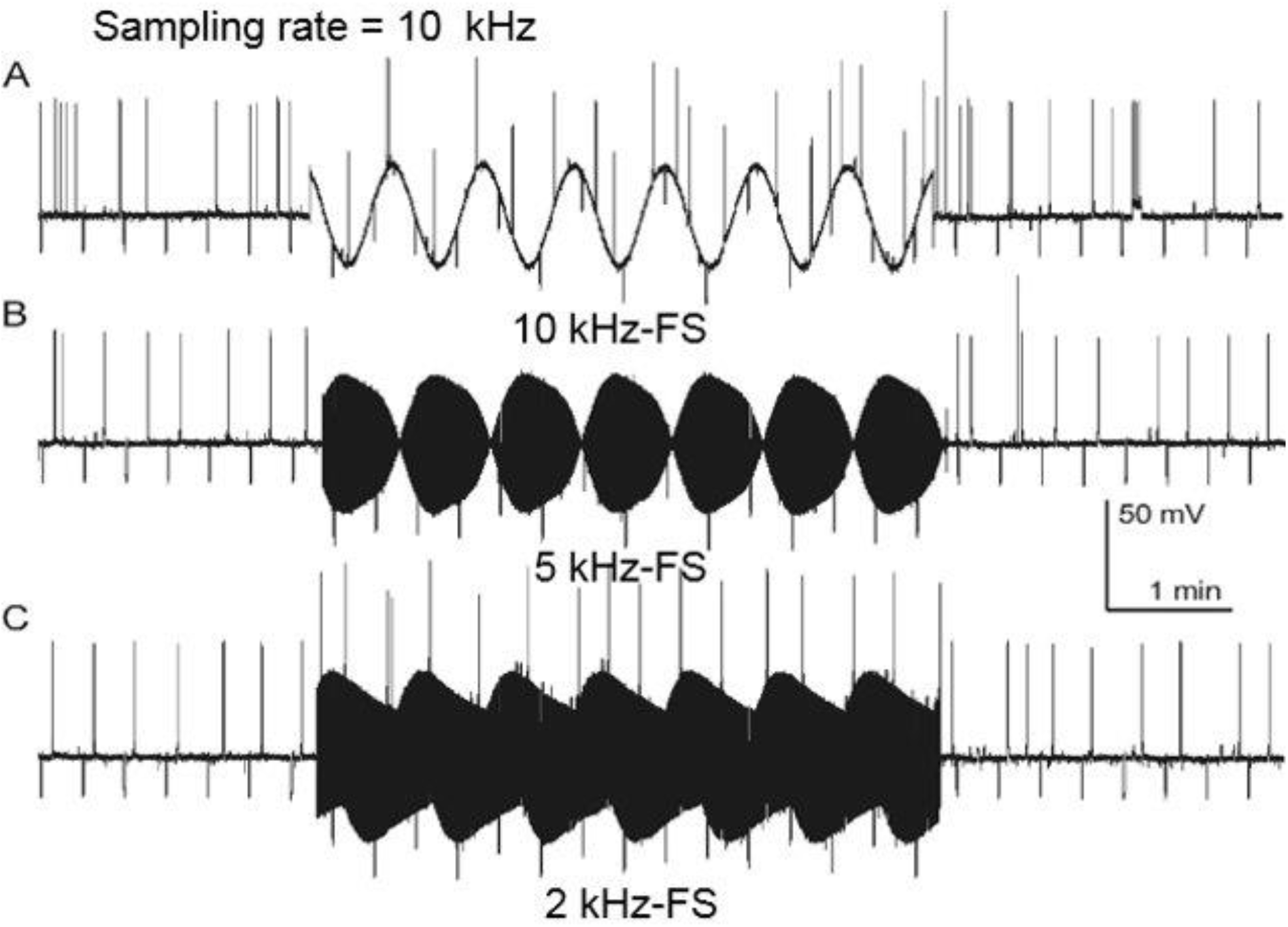
Recording with sampling frequency 10 kHz and different stimulating frequencies. Stimulation with kHz-FS was applied through <u>two bare wires</u> 5 mm long and 5 mm apart located above the slice; the recorded neuron was near the midpoint of the 5×5 mm square between wires. In all three cases <u>stimulation current was 3 mA</u>. No changes in parameter values were observed. **A** - When sampling and stimulating frequencies are the same, the baseline during the stimulation period is smooth, although distorted by the overlapping slow wave. **B** and **C** – inequality of sampling and stimulating frequencies results in high-frequency noise masking the intracellular signal. In *A* testing for Rh continued during 10kHz-FS showing no change. In *B* the neuron was not tested with depolarizing pulses during 5kHz-FS. In *C* it was tested for Rh and generated spikes in response to the same amplitude of depolarizing pulses as it was before and after exposure to 2 kHz-FS.

**Fig. 3.**
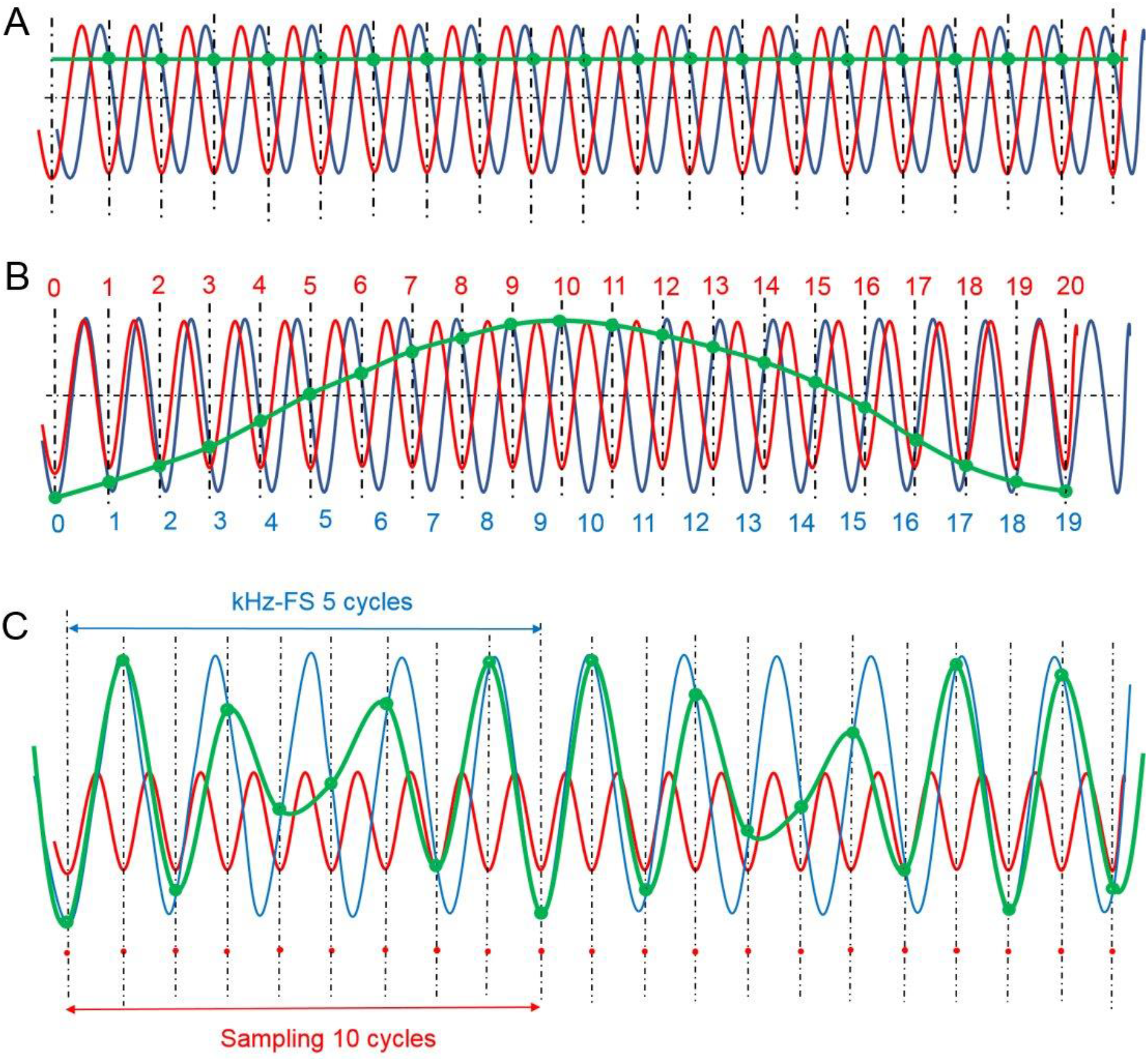
Diagram illustrating interactions between stimulating kHz-FS and sampling (SmF) frequencies. Data acquisition (sampling) of Vm overlapped by kHz-FS (blue lines) occurs at each negative peak of the red line which stands for the sampling frequency. Moments of sampling are indicated by vertical dotted lines and by green circles at their intersection with the intracellular signal. The recorded signal is depicted as a green line connecting sampling points. **A** – The ideal case of equality between SmF and kHz-FS. An arbitrary phase shift between these frequencies results in a positive or negative shift of the Vm baseline; a phase shift equal to a quarter of an oscillation period would result in genuine undistorted values of Vm. **B** –Inequality of the two frequencies, where 20 samples were taken during 19 stimulating oscillations, results in recording of a smooth slow wave overlapping the Vm. Assuming that SmF=10kHz, the slow wave lasts 0.1ms x 20 = 2.0 ms. In our recordings with SmF=10 kHz the slow wave was ∼ 100 s long, i.e. frequency difference was a loss of one cycle per one million (100,000ms/0.1ms=1,000,000). Thus, in our setup frequency difference was one missing or additional kHz-oscillation per a million in SmF relative to kHz-FS. However, it was enough to produce a high amplitude slow oscillation. **C** – A significant discrepancy between SmF and kHz-FS results in recording of a high frequency noise of an alternating amplitude.

**Fig. 4.**
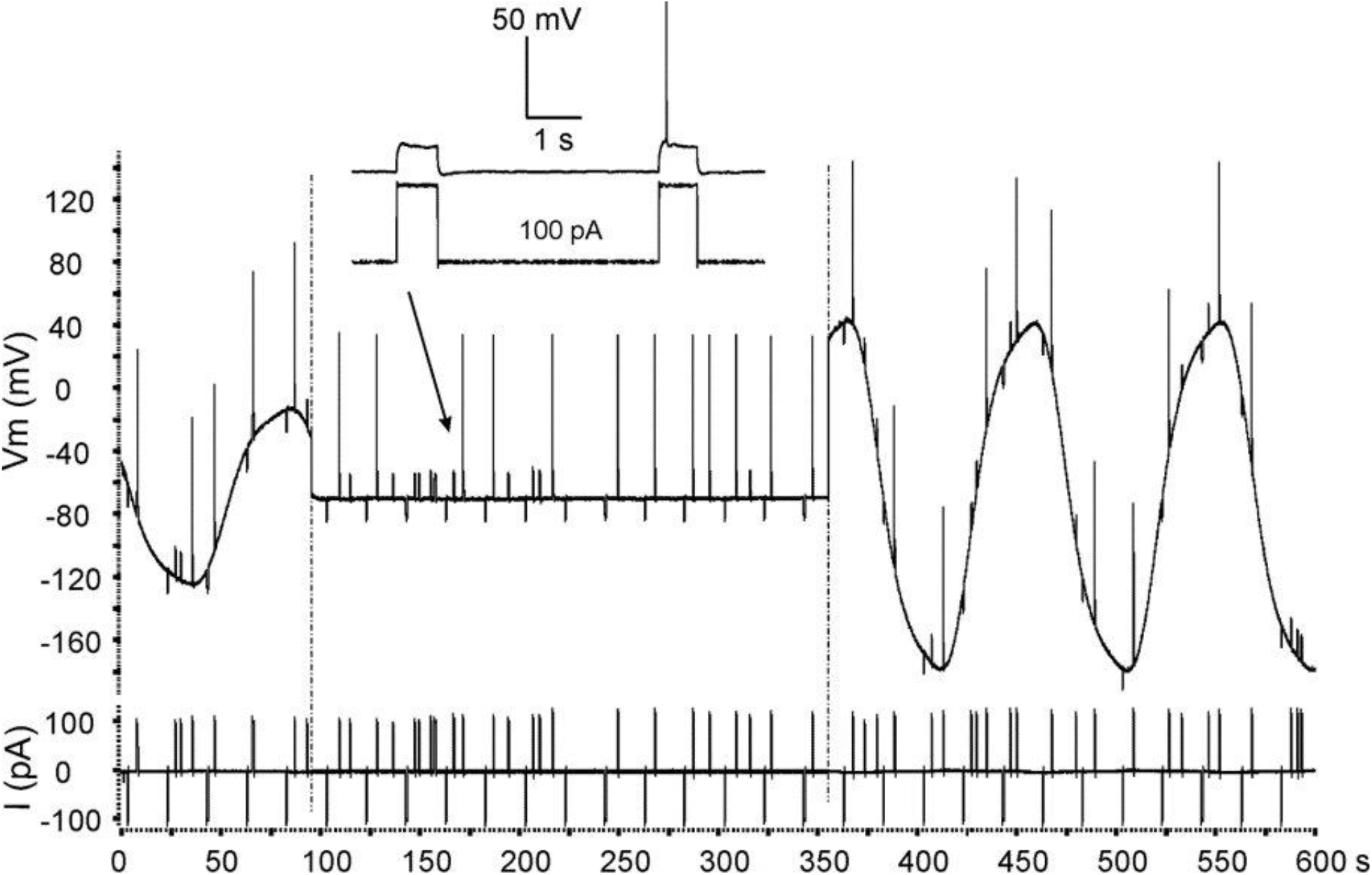
A two-channel 10 min long sweep with intracellular membrane potential (upper trace) and injected currents (lower trace). Regular 500 ms long current injections were made to test Rin (−100pA pulse every 20 s) and Rh (positive current pulses of varying amplitude to adjust for the Rh magnitude). Application of the kHz-FS was always accompanied by occurrence of a slow oscillation superimposed on the intracellular recording. Abrupt ending and beginning of the kHz-FS are marked correspondingly by left and right dashed vertical lines. Therefore, the moments of kHz-FS switching off and on were always distinctively separated from fragments without stimulation. Note equal periods but different amplitudes of slow oscillations. On the left – 5 kHz-FS with 0.5 mA; on the right – 5 kHz-FS with 1 mA. The inset shows two enlarged pulses of just threshold amplitude in response to injections of +100 pA; the arrow indicates these pulses in the sweep; there is a failure in response to the first pulse and occurrence of a single spike in response to the second pulse.

**Fig. 5.**
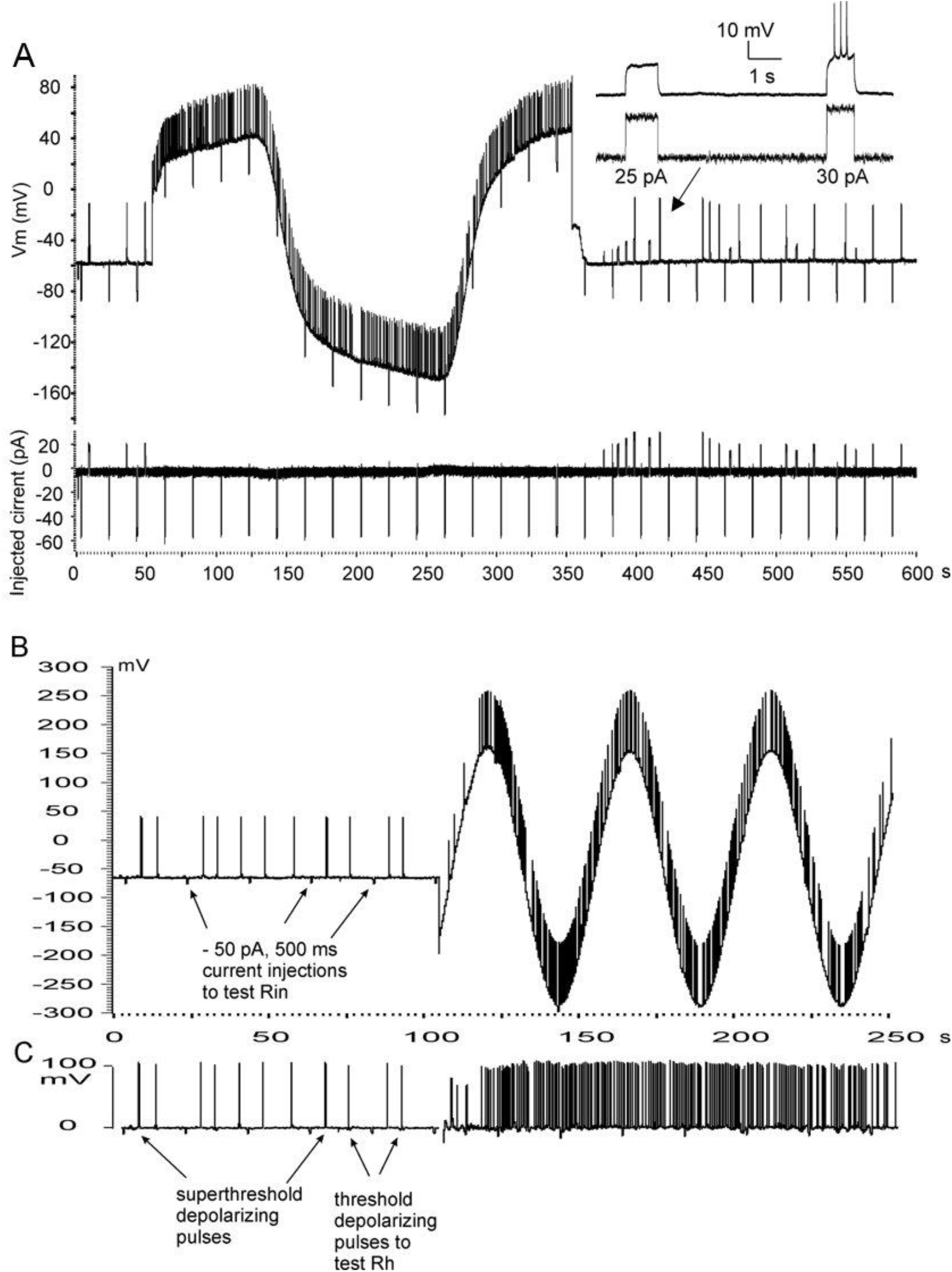
Method for recording neuronal properties during kHz-FS. **A** - recording of a silent neuron responding to intracellular injection of depolarizing current with one or more spikes (the first 50 s on the left). Immediately after the 2 kHz-FS (0.5 mA) was turned on (right half of the trace) the intracellular recording exhibited a large amplitude slow wave along with or combined with the Vm trace. Spontaneous firing of a previously silent neuron shows that the neuron became spontaneously active immediately after the beginning kHz-field and continued firing throughout the rest of the stimulation period. Immediately after cessation of 2 kHz-FS spontaneous firing stopped indicating a decrease of neuronal excitability compared to stimulation period. Furthermore, incremental increase of positive currents injected into the cell showed a transient increase of Rh for ∼100 s after 2kHz-FS was stopped. **B** – A silent neuron tested for Rin with negative current pulses and for Rh with positive current pulses. Amplitudes of positive pulses were varied up and down to adjust for a single spike generation. Turning on 10 kHz-FS (1 mA) initiated spontaneous firing with ∼ 20 s delay. Spontaneous firing persisted with adaptation throughout period of stimulation. In both A and B the changes in neuronal excitability during kHz-FS are obvious despite distortion of traces by slow waves. However, a quantitative assessment of cell properties using the distorted trace is difficult. **C** - To overcome this obstacle we subtracted the modulating wave from the original recording in B. In the resulting trace the Vm baseline became zero. Thus, measurement of resting Vm during kHz stimulation was still impossible, but Rin, Rh and spontaneous firing rate could be measured.

The testing strategy was standardized after a series of pilot experiments where we found that at a fixed stimulating current the strongest response occurred during 2kHz-FS and the weakest response during 10 kHz-FS. Therefore, if a few frequencies at a fixed current amplitude were tested, we started from 10 kHz-FS, then applied 5 kHz-FS and finally 2 kHz-FS. On the other hand, when we tested a range of stimulus amplitudes for a given frequency, we started from the lowest current intensity capable of eliciting response (0.5 mA) with an incremental current increase. Repeatability of the kHz-FS action was tested in some neurons by a series of identical stimuli (Fig.6: 10 kHz 2mA; Fig.7: 5 kHz 1 mA; Fig.8: 2 kHz 1 mA). In each slice only one neuron was recorded. Therefore, previous uncontrolled kHz-field exposures were excluded. Since in this study data points were collected with 20 s intervals, we cannot differentiate whether a parameter was rapid in onset or it occurred at short latency. In most cases cell parameters returned to pre-train values also within 40-60 sec. However, due to standardized data point collection it might be shorter than 20 seconds.

**Fig. 6.**
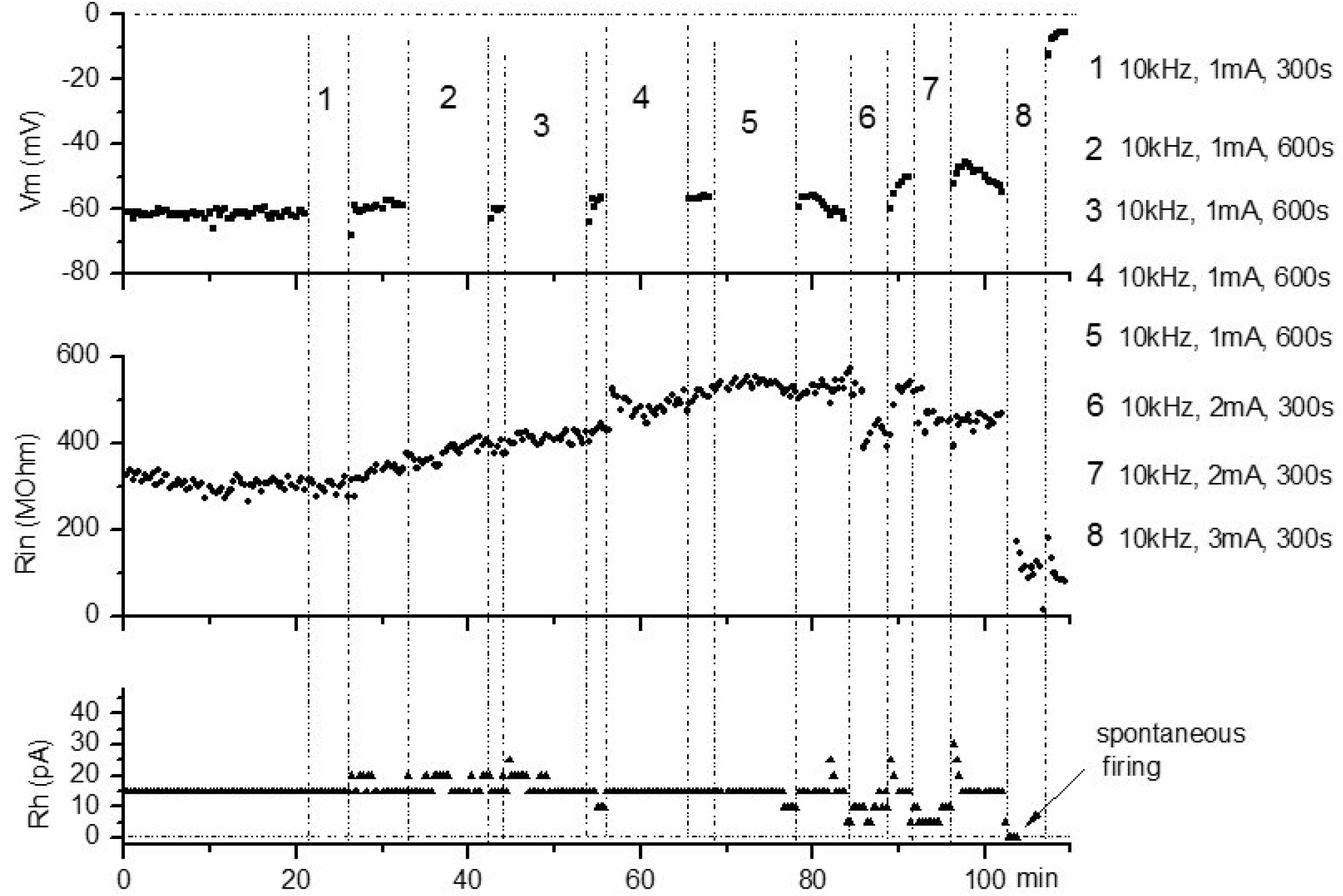
Stepwise increase of 10kHz-FS stimulating current amplitude altered the properties of a neuron. Exposure to five consecutive sessions of 10kHz-FS at 1 mA (##1-5, total 45 min) did not affect the neuron, but an increase of current amplitude from 1 mA to 2 mA decreased input resistance (Rin) and rheobase (Rh) without changing membrane potential (Vm) (#6 and #7). Further increase of current amplitude to 3 mA caused immediate neuronal damage and/or irreversible changes (#8). Monotonic increase of Rin from 25 s to 75 s does not show an accumulation of effects; therefore, it is considered uncorrelated with kHz-FS exposure.

**Fig. 7.**
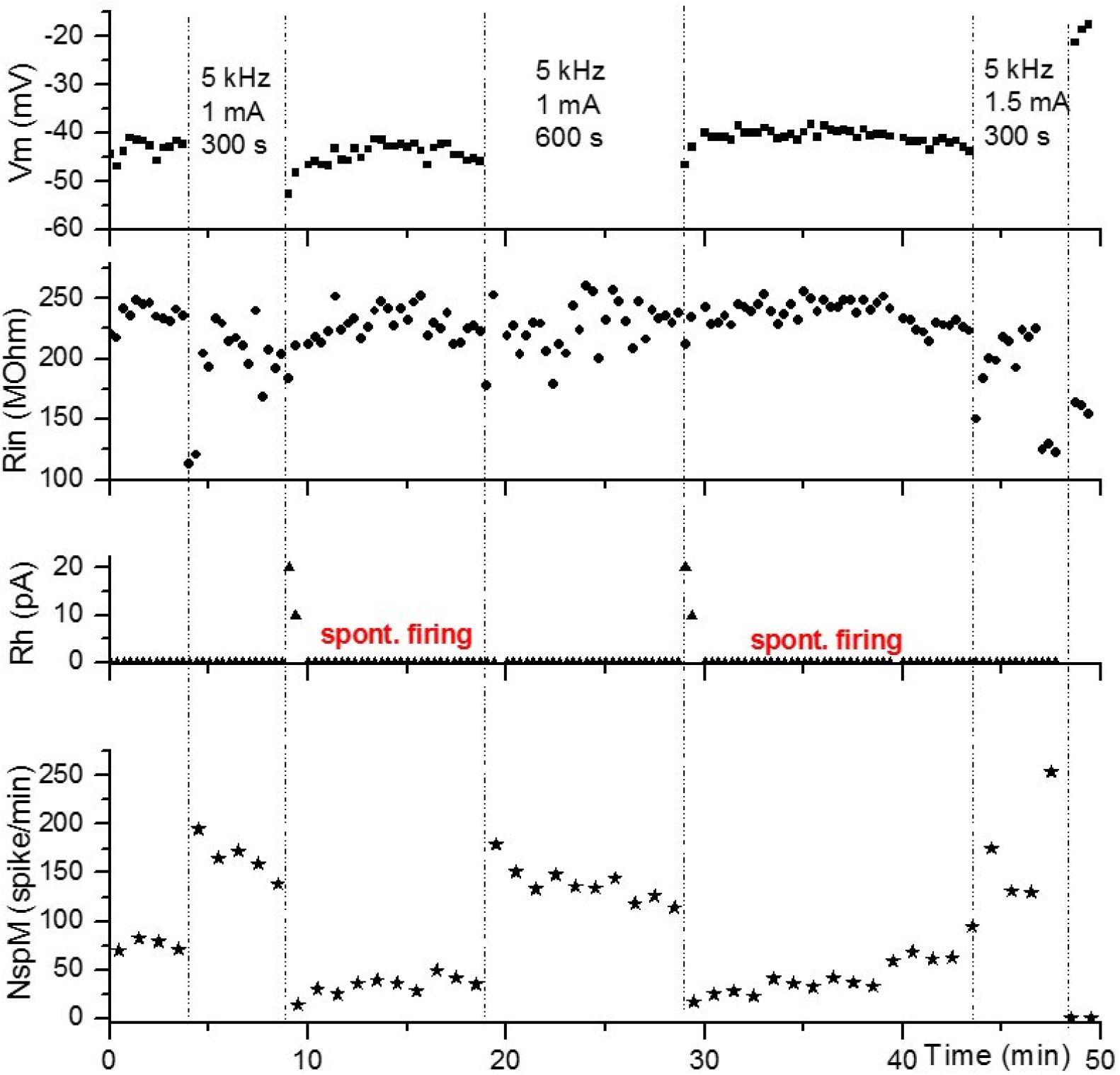
Spontaneously firing neuron did not show changes in Vm, Rin and Rh either during or after 5kHz-FS at 1 mA. However, the firing rate exhibited a rapidly reversible increase during external kHz field stimulation (lower plot). An increase of stimulating current to 1.5 mA led to apparent cell damage and irreversible depolarization at the end of 5-min field application.

**Fig. 8.**
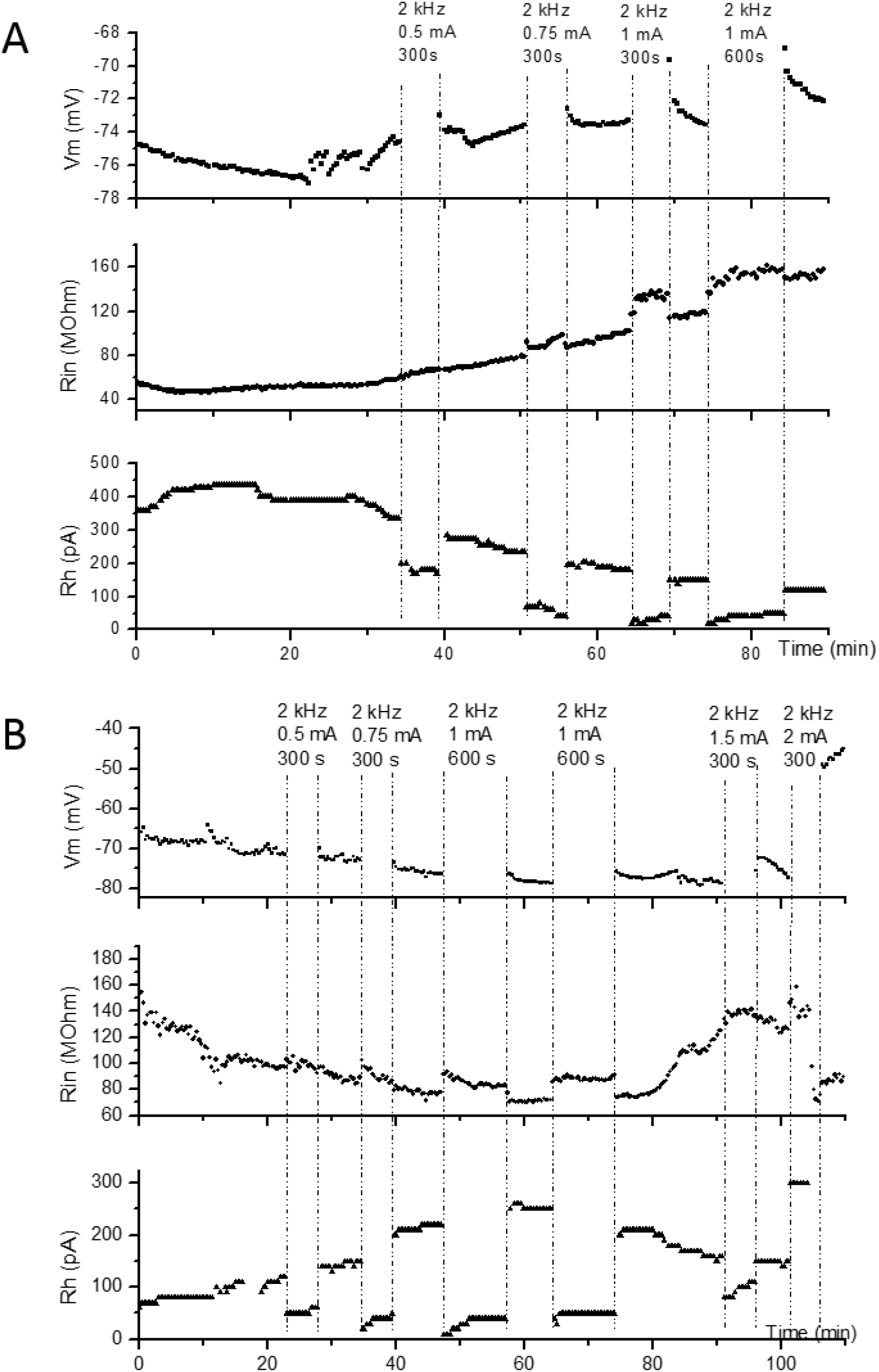
Effects of increasing amplitude of 2kHz-FS on membrane characteristics of two neurons. **A** – 2 kHz-FS induced increases of Rin and decreases in Rh correlated with stimulus intensities. This cell showed an aftereffect only in Vm: where transient increases of Vm after stimulation imply that depolarization occurred during exposure to the external biphasic field stimulation. **B** –This cell showed less prominent Vm tails to the same sequence of stepwise increases in stimulating currents (0.5 mA, 0.75 mA and 1 mA). Increases of Rin and decreases of Rh during stimulation correlated with current amplitudes.

To monitor input resistance (Rin) a 500 ms pulse of negative current was injected via the microelectrode every 20 sec. Rin was calculated by dividing amplitude of the maximal voltage deflection within the first 200 ms (I_h_ sag) by the magnitude of injected current (usually 50 or 100 pA). To monitor rheobase (Rh) positive brief current steps reaching the spike threshold were regularly applied. This minimal injected current was accepted as the value of rheobase. Resting membrane potential (Vm) was measured in front of every Rin-testing negative step only in intervals between kHz-FS. Spontaneous firing occasionally recorded during experiments was accessed by calculated firing rate during and between kHz-FS. Spontaneous firing rate (SpFR) was calculated as number of spikes per minute (not per second) for convenience of placing data points on graphs of long recordings. Statistically significant changes of parameters were assessed using pair-sample T-test (p<0.05). Since most of kHz-FS applications lasted 300 sec, a data fragment representing stimulation period contained 15 data samples taken every 20 s. Therefore, as a control for a given stimulation period, 15 points of the baseline preceding the start of stimulation were taken to be compared with the first 15 data points during stimulation.

## Results

The main finding of this work is the occurrence of changes in neuronal membrane properties and neuronal activity during an exposure to biphasic electrical fields of kHz frequencies. In transverse brain slices submerged in the ACSF, neurons of the hippocampal CA1 subfield were patched blindly and recorded in the whole-cell current-clamp mode for as long as they maintained resting membrane potentials below -40 mV and were able to fire spikes. Recordings for more than one hour were obtained from 27 neurons of *stratum pyramidale* while they generated action potentials in response to positive current injections or fired spontaneously. Slowly developing changes of neuronal properties due to prolonged recording could be distinguished from that caused by kHz-FS based on several characteristics. First, monotonic trend in Vm, Rin or Rh changes was often observed throughout 1-2 hours long control recording (not shown) similar to trends interpolated from inter-kHz-FS fragments, whereas changes elicited by kHz-FS were always abrupt (Fig.6, Fig.8, Fig.9). The former probably results from a few factors intrinsic to whole-cell method: cell dialysis with the intra-pipette solution, cell swelling and increasing leakage of the membrane. Second, in our experiments slow gradual parameter changes often started before the first kHz-FS train (Fig.8). Third, on plots the post-kHz-FS values of parameters were in most cases predicted by extrapolation. This indicated independence of a gradual monotonic change from exposure to electrical stimulation. In fact, we saw extremely rare (in 3 of 27 neurons) reproducible post-kHz-FS transient depolarization (Fig.6A and 8A) and (1 of 27 cells) a brief post-kHz-FS hyperpolarization (Fig. 7). In 4 of 27 neurons we have observed post-kHz-FS shifts of parameter values which could last until the next application of kHz-train (Fig. 9B, Vm after 2kHz 0.5 mA); this cannot be interpreted as actual persistent post-stimulating effects because they were not reproducible and hard to explain. Two cells showed steadily depolarized Vm after the last kHz-FS (Fig.9A). One cell showed a stable shift of Vm after 2kHz-FS at 0.5 mA but after the following 2kHz-FS at a higher intensity of 1.0 mA the Vm returned to the approximated baseline (Fig.9B); this cannot be interpreted as a persistent change. Another cell showed Vm decrease and Rin increase after the first cycle of 10 kHz-FS at 1 mA but did not respond with any parameter changes to the second exposure to 10 kHz at 1 mA; i.e. its response was not reproduced. Therefore, we concluded that no reproducible persistent aftereffects were observed. In cases when obvious aftereffects occurred, (a) they were caused by the highest amplitude of biphasic currents applied to the neuron, (b) showed strong depolarization and loss of Rin and (c) were irreversible. On the contrary, obvious changes of whole-cell parameter values during kHz-FS were always abrupt, repeatable and dependent on an amplitude and/or frequency of kHz-FS.

**Fig. 9.**
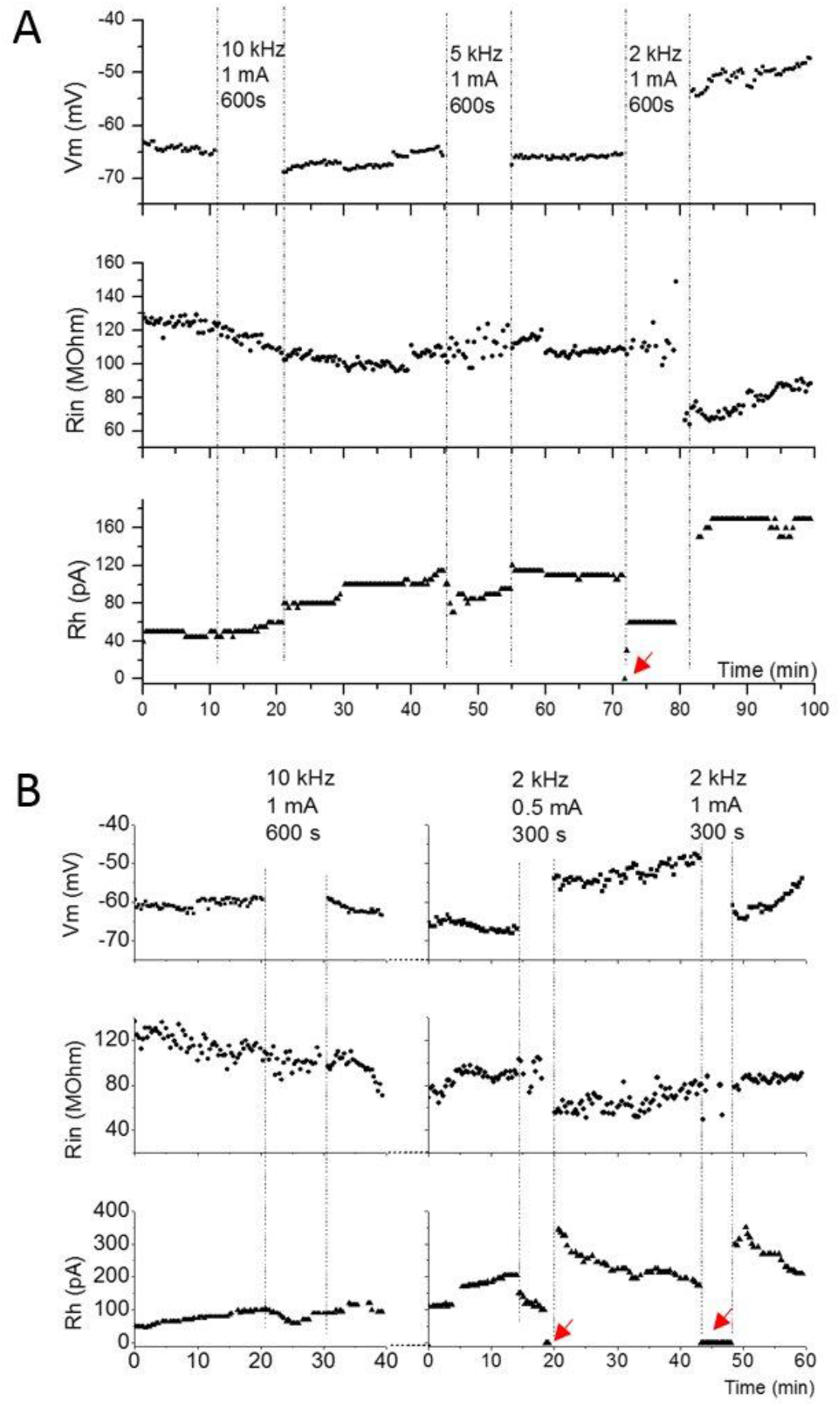
**A** – Application of kHz-FS of the same current amplitude (1mA) but at different frequencies shows no effect at 10 kHz, a small effect at 5 kHz and a stronger effect at 2 kHz. Red arrowhead directed to a point of Rh=0pA indicates transient occurrence of spontaneous firing (not shown) at the beginning of 2 kHz application. **B** – Another example of 10kHz-FS at 1 mA without a noticeable effect (left column of plots) and a well pronounced effect of 2kHz-FS (right column). In this cell 2kHz-FS of the lower intensity (0.5 mA) caused instability of Rin and a significant drop of Rh with a few spontaneous spikes (red arrowhead) followed by a sharp rise lasting for ∼10 min. Aftereffects of 2 kHz-FS at 1 mA included a persistent increase of Vm, a transient decrease of Rin lasting for ∼15 min and a transient increase of Rh lasting for ∼10 min. Recordings with 2 kHz sampling rate (right) were made 5 min after recordings with 10kHz sampling rate. The 5 min gap was needed to switch between the recording protocols. Aftereffect of 2kHz-FS at 1 mA was somewhat different from that at 0.5mA: Vm transiently returned to a control level, Rin did not change, whereas Rh exhibited the same transient increase for 10 min. Red arrows at Rh=0 designate spontaneous firing.

A shift in excitability of a neuron was determined either by a change in magnitude of Rh or by a change of spontaneous firing rate (SpFR). When spontaneous firing was present, Rh was assigned to be zero and positive current injections were cancelled (Fig.7). Any persistent and repeatable change of Rin, Rh or SpFR during kHz-FS was called a response. A transient or persistent clearly observed shift of post-kHz-FS values of Vm, Rin or Rh (compared to pre-FS values) was called an aftereffect. Response of a neuron was considered excitatory if kHz-FS decreased Rh or increased SpFR. Since moments of spike generation during exposure to kHz-FS were arbitrary, and firing might be irregular or adapting, we considered this firing analogous to a genuine spontaneous firing. Therefore, we hypothesize that spikes were elicited by a general superthreshold depolarization but not by regular field oscillations or current injections. Spikes elicited by positive currents injected for Rh testing were excluded from SpFR analysis. In our experiments spontaneous firing could either emerge in silent cells due to decrease of Rh to zero, or it could become more intense during kHz-FS if it existed before a stimulating train. Decrease or loss of spontaneous firing would be considered as decrease of excitability.

We found that effective amplitudes of external field stimulation to induce an excitatory response during stimulation were proportionally correlated with its frequency. Currents as low as 0.5 mA almost always elicited excitatory effects at 2 kHz (8 of 9 cells, Figs. 8), but rarely at 5 kHz (in 1 of 10 cells), and never at 10 kHz. Currents of 1 mA amplitude were rarely shown to be effective at 5kHz-FS (2 cells of 9, Fig.7, Fig. 9A) or at 10kHz-FS (3 cases of 11). However, responses to 2 mA at 5kHz-FS were elicited in most cells (5 of 7). At the same time 2 mA represented only a threshold value at 10kHz-FS (Fig.6). Similarly, at the same stimulating frequency an increase of stimulus amplitude elicited more prominent responses of a neuron (Fig.8).

Dependency of responses on the amplitude and frequency of kHz-FS is summarized in Figs. 11 and 12. Overall, the largest percentage of neurons responding to stimulation intensity lower than 2 mA occurred during 2kHz-FS (100%), whereas this percentage was the lowest at 10kHz-FS (18%) and intermediate at 5kHz-FS (45%). Rin either did not change or decreased during kHz stimulation and returned to pre-stimulation values after that. This probably suggests a reversible leakage increase or activation of the Na^+^-persistent current through the cellular membrane caused by the exposure to the kHz field. In all observed cases Rh only decreased and SpFR (when expressed) only increased indicating an exclusively excitatory effect of kHz-FS on neurons. It seems that the increased level of excitation could be maintained or repeatedly obtained with a certain current amplitude at a given stimulation frequency (Fig.7: 5kHz, 1mA; Fig.8: 2 kHz, 1mA). In none of the cases did we observe an increase of Rh or a decrease of SpFR during stimulation. Therefore, external kHz electric fields produce only excitatory but not inhibitory effects on hippocampal pyramidal neurons during stimulation. Random and rare spontaneous IPSPs did not change their rate of occurrence during kHz-FS, indicating that GABA-ergic inhibitory neurons synapsing with a recorded cell were not excited. Occurrence of spontaneous EPSPs also was not changed by kHz-FS indicating that synaptic input from CA3 subfield was not affected.

Finally, we wanted to test whether effects of kHz-FS represent a direct action of the high-frequency external field on the recorded pyramidal cell or mediated by targeting the neuronal network and altering the synaptic input to the pyramidal cell. Therefore, a cocktail of inhibitors for glutamatergic AMPA and NMDA receptors (CNQX 10 mM and D-AP5 50 mM correspondingly) was added to the bath to block glutamatergic synaptic transmission (3 cells tested). Fig.10 demonstrates that synaptic block did not eliminate the excitatory effect of 2kHz-FS at 1.5 mA and even did not change the firing rate during field application (compare #4 – pre-block, #5 synaptic block and #6 – washout).

**Fig. 10.**
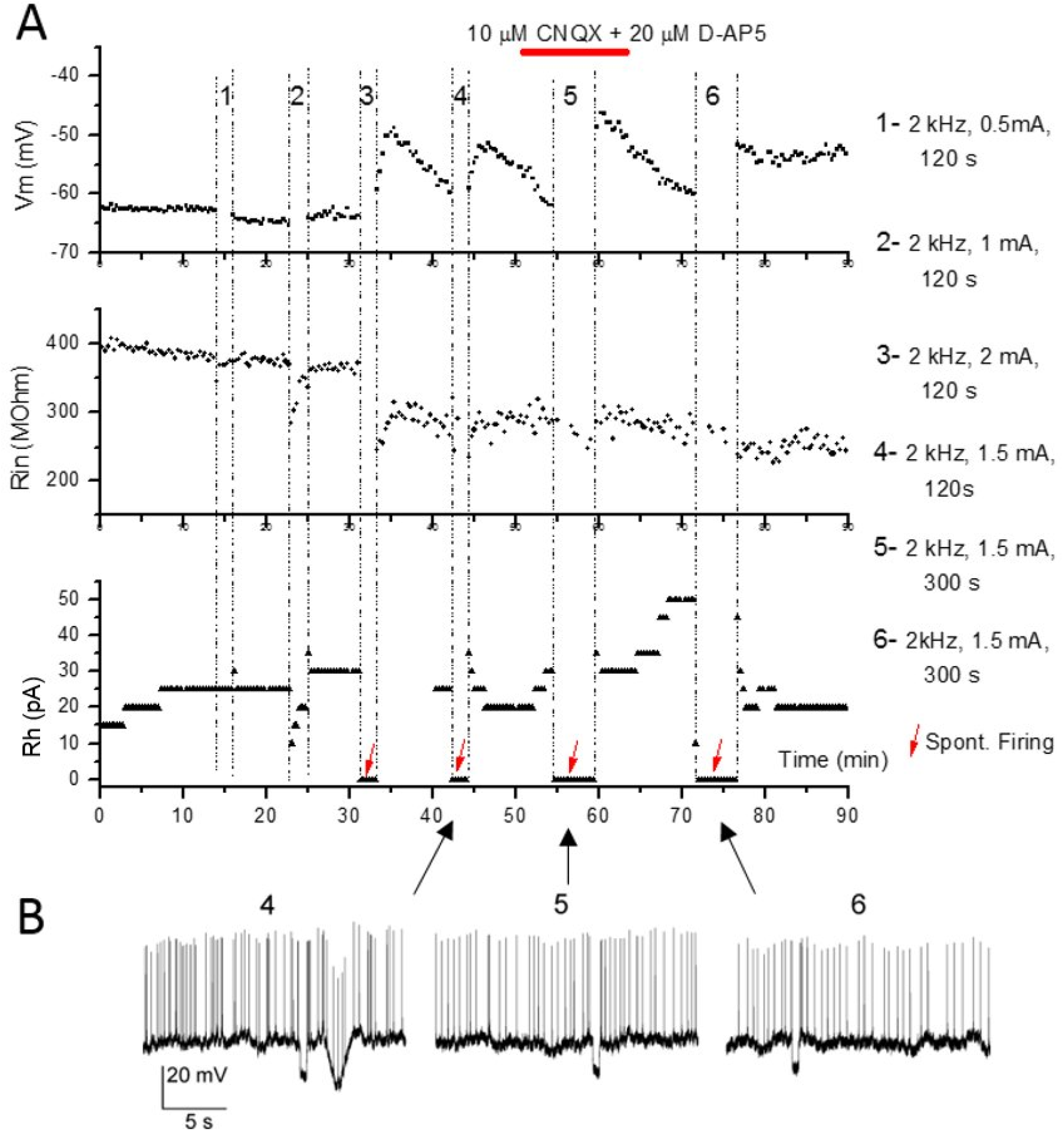
**A** – Successive applications of 2 kHz electric field with incrementally increasing currents induced almost no response to 0.5 mA (1), a weak transient decrease of Rin and Rh at 1mA (2) and a superthreshold excitation of the neuron during 2kHz-FS at 2 mA followed by irreversible decrease of Rin (3). Stimulation with an intermediate intensity of 1.5 mA resulted in similar responses before (4), during (5) and after bath application of synaptic blockade of glutamatergic synaptic transmission (6). **B** – Firing patterns elicited by 2kHz-FS at 1.5 mA before (4), during blockade of AMPA-R and NMDA-R (5) and after washing out the drugs in ACSF (6). Similar frequency of firing indicates a direct effect of the kHz-field on the recorded cell with no or a negligible impact of excitatory synaptic inputs from adjacent or remote neurons.

**Fig. 11.**
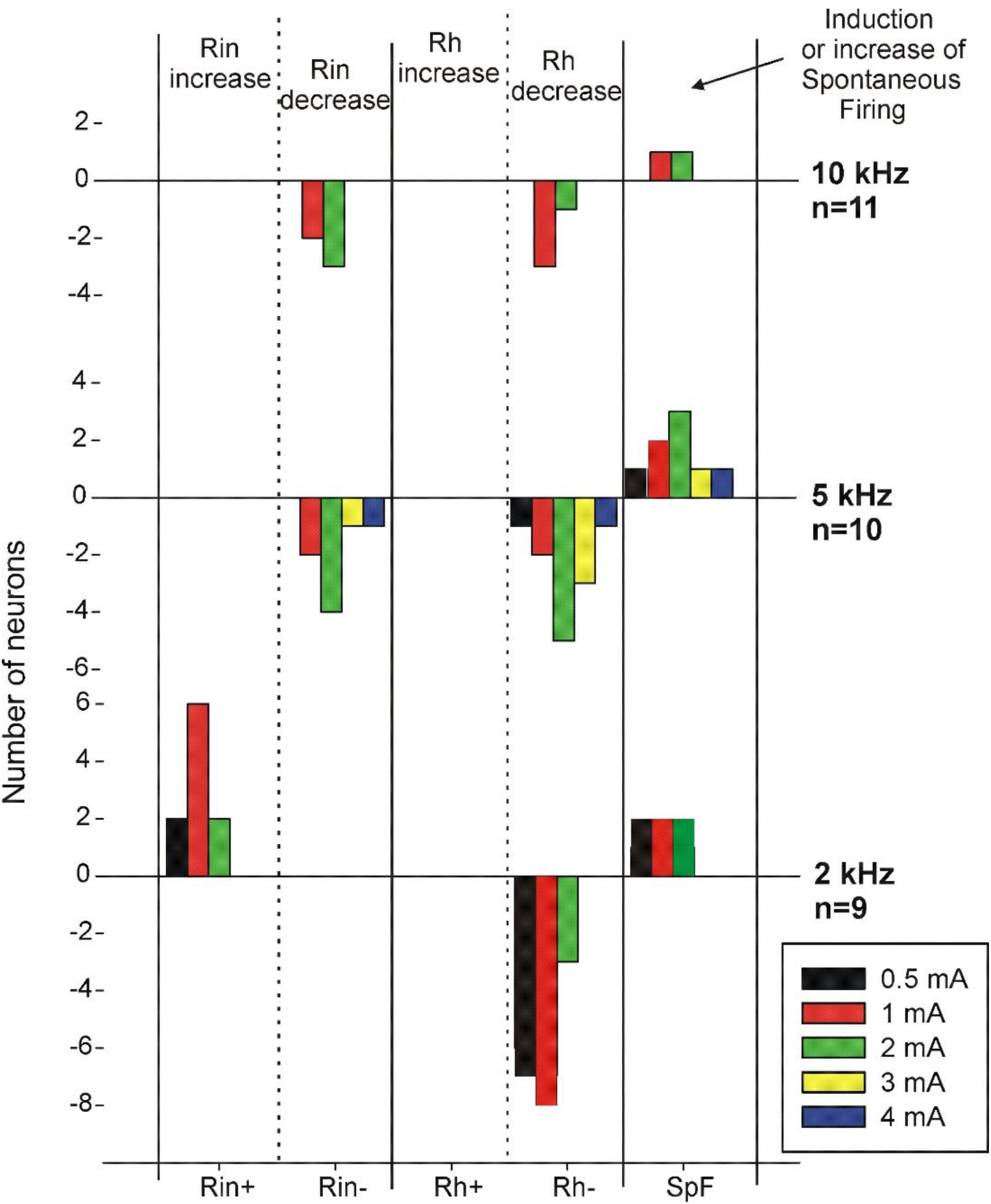
The colored bar graph illustrating direction of parameter changes and their sensitivity to kHz-Fs. Positive numbers correspond to increased parameter values and negative numbers correspond to decreased values. Rh was the most sensitive parameter showing only a decrease at all stimulus frequencies and intensities. 10 kHz-FS was the least efficient and 2 kHz-FS was the most efficient in eliciting responses. Rin showed a prominent frequency dependence only increasing during 2kHz-FS regardless of a current amplitude but decreasing during 10kHz-FS and 5kHz-FS. Total number of neurons tested with each kHz frequency is on the right side of the graph. Neurons used for this graph were not all tested with different frequencies, but in all of them stimulation started from the lowest, usually subthreshold intensities. Therefore, the same neuron responding to a few incremental current amplitudes was counted for each of them.

**Fig. 12.**
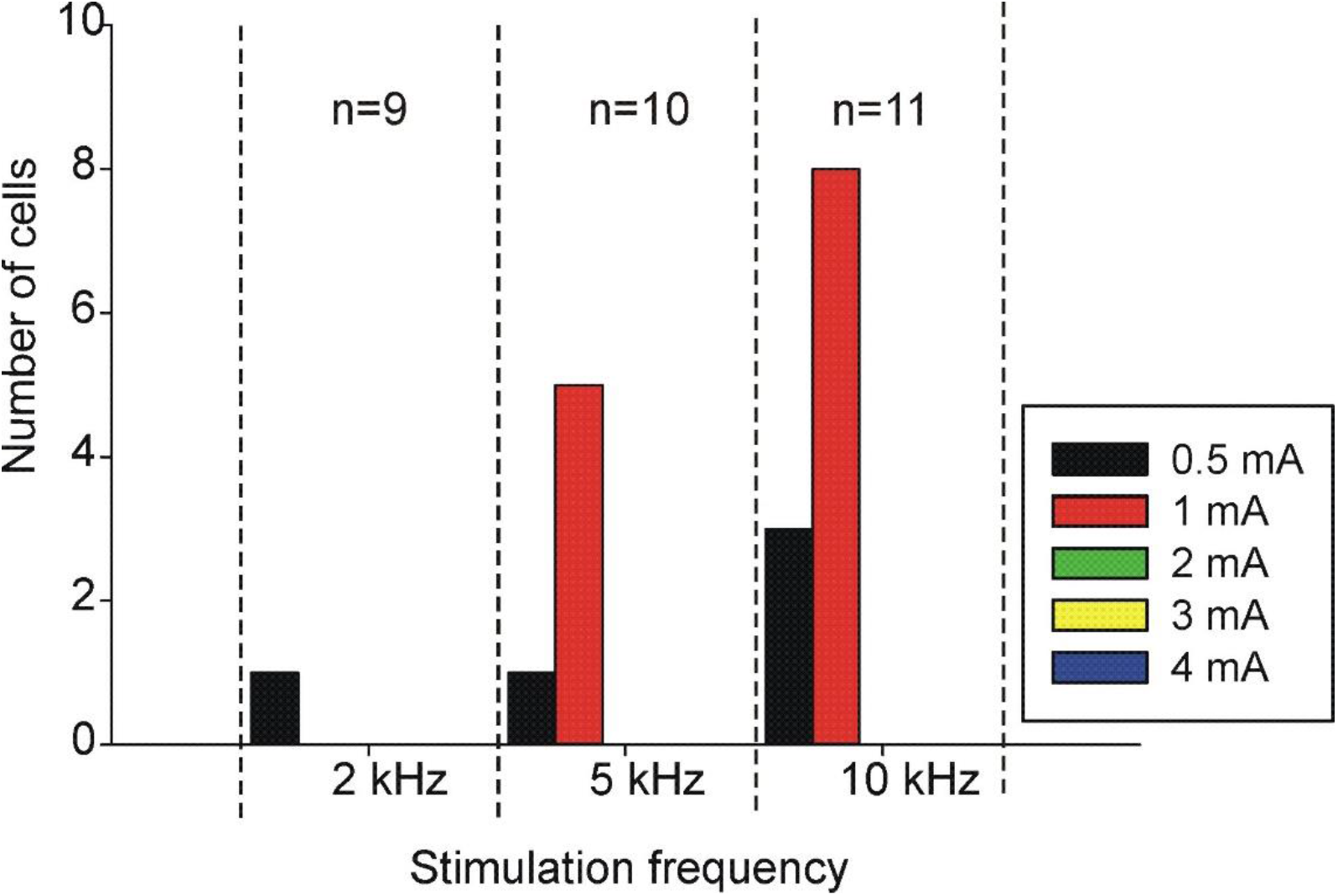
Numbers of cells which showed no changes during kHz-FS at any frequency or intensity of stimulation. At 10 kHz most and at 5 kHz a half of tested neurons did not respond to currents lower than 2 mA. At the same time only 1 of 7 cells did not respond to 0.5 mA 2kHz-FS indicating that lower kHz-frequencies have a higher probability of inducing excitatory effects. Total number of neurons tested with each kHz frequency is indicated above vertical bars within each column. In this figure 8 of 11 cells did not respond to FS<2mA, whereas other 3 of 11 did respond to 2mA. These three cells are included in Fig. 11. All three cells changed Rin and Rh, and one of them started spontaneous firing. Only 3 cells were tested with 0.5mA at 10 kHz with no response. Therefore, 0.5mA was considered the subthreshold stimulus intensity at 10kHz. For other cells 10 kHz tests started from 1 mA, where 8 of 11 cells failed.

Our data with large variability of cell properties and with uncontrolled changes in the course of long recording often did not allow comparison for significant differences between pre-stimulus and post-stimulus fragments due to trends usually present in recordings. For example, in Fig.8 Rin has a rising trend whereas Rh has a decaying trend. Because of this very often pre-kHz-FS fragments were significantly different from corresponding post-kHz-FS fragments. However, this difference was due to a gradual trend, and therefore it could not be considered as an after-effect. Hence, we could test for significant differences only pairs of pre- and post-stimulation fragments where a trend was minimal or absent, and the baseline was flat. For example, in Fig.9B exposure to 2 kHz 0.5 mA led to a significant rise of Vm, significant but transient decrease of Rin and increase of Rh. On the other hand, all changes of parameter values during stimulation represented abrupt shifts and differences were significant when the slope of a trend was subtracted (overlapping trend was often observed throughout the stimulation period). For instance, in Fig.9 the tilted parameter fragments during stimulation indicate presence of a conserved trend. Nevertheless, comparison of a pre-stimulus fragment with a shifted corresponding fragment during stimulation showed significant differences regardless of a trend.

## Discussion

Thus, qualitative assessment of data combined with testing for significant changes during stimulation leads to a few unequivocal conclusions. First, exposure of neurons to the kHz-frequency biphasic electric fields does induce an increase in neuronal excitability evident as a reduction of the rheobase and an increase of average spontaneous firing rate. Second, expression of kHz-field effects is negatively correlated with frequency of the field stimulation and positively correlated with the current amplitude. In this study we did not identify exact threshold amplitudes of kHz frequencies but, roughly, superthreshold values of currents were: 0.5 mA at 2kHz-FS, 1-1.5 mA at 5kHz and >2mA at 10 kHz. Third, long exposure to subthreshold kHz-FS for up to an hour (Fig.6, 10 kHz, 1 mA, 45 min) does not induce detectable and persistent changes in baselines of measured parameters. Fourth, the best indicators of kHz field effects were Rh and the rate of spontaneous firing. Fifth, in most cases the effects were reversable within seconds after cancellation of stimulating kHz currents.

The intrinsic low-pass filtering of the neuronal membrane explains limited sensitivity of neurons and neuronal networks to kHz electric fields [20-22, 29-32], but it does not exclude an existence of specific biophysical mechanisms responsible for effects of kHz-FS. To investigate changes and processes occurring in a neuron while it responds to external kHz field, one needs to develop a reliable model producing such responses on a regular basis. We developed such a model and in this publication we first demonstrate an unequivocal excitatory effect of kHz-FS on neurons. It was found that all selected parameters characterizing neuronal excitability do change in a threshold-dependent manner. A response threshold increases with increase of stimulating frequency. It is possible, although not tested here, that response thresholds differ for different neurons and depend on their initial states, intrinsic properties, and external conditions.

An acute hippocampal slice has been used previously by several investigators to study effects of a range of stimulation waveforms and intensities [23, 33]. Structure, connectivity and properties of neurons in the CA1 region have been thoroughly studied for the past decades. In CA1 region excitatory synaptic contacts between principal neurons are extremely rare [34], although adjacent neurons can be connected through gap-junctions [35, 36]. The major synaptic input to apical dendrites of CA1 neurons comes from CA3 subfield. Therefore, if only CA1 subfield is exposed to the external kHz-FS, even without blockade of glutamatergic transmission they may be considered isolated from excitatory synaptic inputs. At the same time, pyramidal neurons of CA1 subfield receive an abundance of inhibitory inputs from local interneurons [6, 37]. Basket, bistratified and axo-axonic cells are parvalbumin (PV) expressing GABA-ergic neurons whose somata are intermingled with pyramidal cell bodies in the pyramidal cell layer of CA1 [38, 39]. They can be reliably differentiated from pyramidal cells by a characteristically narrow spike and a very deep after-hyperpolarization (AHP). Each PV-interneuron targets hundreds of pyramidal cells. Therefore, being activated they can induce a powerful hyperpolarization in pyramidal neurons suppressing their excitability and spontaneous firing. Thus, if local GABA-ergic interneurons were excited by the kHz-FS they would feed-forwardly inhibit pyramidal cells during kHz-FS but we did not observe lowering of pyramidal cell activity or increased density of IPSPs in recorded traces. Lack of inhibition in pyramidal cells may be due to non-overlapping levels of sensitivity to the external kHz field in different neuron types. One can speculate that a kHz-FS induces suppression of activity in the inhibitory network, but it needs to be tested directly in pyramid-interneuron paired recordings. Another option is that an average threshold of CA1 inhibitory neurons is either much higher than that of principal neurons, or much lower. In this case at stimulus intensities eliciting response in pyramidal cells, the inhibitory interneurons would be inactive either due to insufficient activation or vice versa due to overexcitation with a low field intensity following depolarizing block. This hypothesis is feasible in the light of recent finding that excitatory and inhibitory neurons of the dorsal horn respond differently to kilohertz frequency spinal cord stimulation [40, 41]. The authors found that in the *in vivo* and *in vitro* configurations the low-intensity 10kHz (but not 1kHz or 5kHz) spinal cord stimulation selectively activates inhibitory neurons without activating excitatory neurons and ascending axons of the dorsal column. Thus, an additional series of experiments with simultaneous paired recordings from excitatory and inhibitory neurons is necessary to reveal possible divergence between responses of different cell types in the hippocampus and neocortex.

To achieve a nerve conduction block the frequency and amplitude of the kHz-FS need to be in certain relations, namely, they should be positively correlated. Block threshold amplitudes have been shown to increase linearly as a function of stimulus frequency in fast (myelinated) fibers; whereas in slow conducting (non-myelinated) fibers correlation was also positive, although with a greater dispersion [25, 42, 43]. In our experiments the lowest average threshold amplitude was found for the lowest stimulating frequency (2 kHz). However, the effect was quite opposite to the case of nerve stimulation. Instead of an excitability decrease the CA1 pyramidal neurons showed unequivocal excitatory responses at all frequencies when stimulation intensities were within the physiological range. We hypothesize that if the mechanism of kHz-FS effect is the same in the nerve fiber and in the pyramidal cell, the nerve might undergo a reversible and lasting depolarizing block without membrane damage. In neurons the soma depolarizes and generates spontaneous firing during exposure to low and moderate kHz-FS intensities, but cannot tolerate high kHz-field intensities and dies presumably due to massive Ca2+ influx and excitotoxicity during prolonged stimulation [44].

However, an existence of different mechanisms of kHz-FS action on a nerve fiber and neuronal soma is also possible. Electrode design, geometry and distance to the axon play a role in the efficacy of conduction block [45, 46]. For bipolar cuff electrodes computational experiments demonstrated a monotonic increase of block threshold and extension of the onset response with increasing distance between electrodes placed along the axon axis [47, 48]. Deep brain stimulation (DBS) applying high-frequency currents to deep brain structures which is now an established treatment for various neurological and psychiatric diseases usually employs a thin plastic shaft with a few circular or semicircular conducting rings along the shaft. The shaft is stereotactically implanted in a targeted structure where selected contacts create a symmetrical or asymmetrical electric field [49]. Configuration and spread of the field in the tissue modulates the physiological effects of treatment. For example, although conduction block can be achieved with monopolar, bipolar and tripolar cuff electrodes, monopolar stimulation shows longer onset responses [48, 50]. In our configuration we attempted to minimize non-uniformity of the electrical field to exclude variations of field intensity for recorded neurons differently positioned relative to stimulating electrodes. This design also provides a possibility to obtain quantitative assessment and comparison of kHz-FS action on different types of neurons.

It seems that minimization of kHz-current shunted through ACSF in experiments with submerged slices is essential for obtaining a measurable neuronal response. When stimulating wire electrodes were placed in the bath on the sides of a slice but mainly were contacting the ACSF, no significant field effect was observed in the experiments of [24] and in our pilot experiments (not shown). Passing stimulating kHz-currents between parallel bare wires 5 mm long and 5 mm apart located above the slice also did not elicit any response at 3 mA (Fig.2), although in the setup used in the current study the kHz-FS of this amplitude was superthreshold at all frequencies. Passing the current predominantly through the tissue, as it takes place in the *in vivo* case, increases chances of obtaining a reasonably well-expressed neuronal response. In the ideal scenario, the field between parallel wires is isopotential and currents flowing through any point between the wires are the same provided the tissue is homogenous and has the same impedance in all points. Although a spatial profile of currents in real tissue is far from being ideal, small shifts of a recorded cell location in the vicinity of the middle of the imaginary rectangle of the field should not critically affect results. It is still to be tested whether distance from the soma to one of the electrodes or orientation of apical dendrites in the field play a role.

More data with different experimental design for kHz field application is to be collected for more detailed description and quantitative assessment of the external field action. Neurons of different structures in the peripheral autonomic and central nervous system may respond differently to same kHz-FS. Distance between bipolar stimulating electrodes may also play a role. Distortion of the applied kHz-field in the brain or spinal cord due to spatial variations in tissue conductivity will make field effects even more complex. Nevertheless, all the complexities can be studied using a standard well controlled model system, and adequate protocols can be designed for clinical bio-stimulation.

## Abbreviations

AHP: after-hyperpolarization
DBS: deep brain stimulation
kHz-FS: kilohertz-frequency field stimulation
ACSF: artificial cerebrospinal fluid
Rin: membrane input resistance
Rh: rheobase
SpFR: spontaneous firing rate
Vm: baseline membrane potential

## Acknowledgements

We thank Dr. Igor Kurnikov for analysis software development. SK developed the experimental design, conducted experiments, analyzed data and prepared draft of the manuscript. WdG and TC discussed results and edited the draft. The work was supported by NIH grants R01DK129194 to SK and NIH/NINDS grant R01NS109198 to CT, and NIH/NIDDK grant R01DK111382 to JB and WdG.

